# Predicting localized affinity of RNA binding proteins to transcripts with convolutional neural networks

**DOI:** 10.1101/2021.06.02.446817

**Authors:** Alexander Kitaygorodsky, Emily Jin, Yufeng Shen

**Author notes:** Correspondence should be addressed to Y.S.

## Abstract

RNA binding proteins (RBPs) are important regulators of transcriptional and post-transcriptional processes. Computational prediction of localized RBP binding affinity with transcripts is important for interpretation of genetic variation, especially variants outside of protein coding region. Here we describe POLARIS (**P**rediction **O**f **L**ocalized **A**ffinity for **R**BPs **I**n **S**equence), a new deep-learning method for achieving fast, site-specific binding affinity predictions of RNA-binding proteins (RBPs) to the transcribed genome. POLARIS has two modules: 1. a convolutional neural network (CNN) to predict overall RBP binding within a region based on transcript sequence content and expression level; 2. a Gradient-weighted Class Activation Mapping (GradCAM) implementation for efficient signal backpropagation to individual sequence positions. We trained the model using enhanced crosslinking and immunoprecipitation (eCLIP) data from ENCODE. POLARIS has good performance with a median AUC ~ 0.96 for 160 RBPs across three different cell lines, substantially higher than selected popular published methods trained and tested on the same data sets. When tested on data from a different cell line with the same RBPs, the overall performance is maintained, supporting the ability of cell-type specific affinity prediction. Finally, the GradCAM module allows the model to identify the informative sites in a region that drive prediction. The localized prediction facilitates interpretation of the results and provides basis for inference of functional impact of noncoding variants.

## Introduction

Recently several studies have shown that post-transcriptional regulation acts as a major link between rare noncoding variants and human disease [1–3]. Post-transcriptional regulation is mediated by interaction of RNA-binding proteins (RBPs) and messenger RNAs. Leveraging RBP-RNA interaction can help to improve the interpretation of noncoding variants in disease studies. However, our current understanding of the impact of genetic variation on RBP-RNA interaction is limited due to the complex nature of RBP binding: although RBP binding is foremost driven by biochemical recognition of specific sequence motifs, many other factors such as RNA secondary structure also contribute. Indeed, most motifs [4] still remain unknown due to limited *in vivo* binding data, variability in RBP motif strength, and a complicated relationship between raw sequence and true biological binding of RBPs: mere presence of a motif alone does not guarantee binding at a genomic site in a particular tissue. Therefore, to fully understand the effects of rare noncoding variants in post-transcriptional regulatory regions, it is first critical to model underlying RBP binding with greater accuracy.

In the past few years, large efforts such as the ENCODE (Encyclopedia of DNA Elements) project [5] have performed eCLIP (enhanced crosslinking and immunoprecipitation) footprinting experiments to capture in vivo binding of RNA binding proteins, allowing us to perform large-scale genomics data analyses [6]. New methods such as BEAPR [7] have leveraged this crosslinking data to better identify allele-specific binding events and prioritize functional genetic variants that likely mediate post-transcriptional regulation. Additionally, significant advancements in deep learning have provided us with powerful computational tools to solve complex pattern recognition problems related to scanning of binding motifs. Current transcription factor (TF) and RBP binding models such as the DeepSEA/SeqWeaver [8, 9] and DanQ [10] models have seen initial success in modeling binding to sequence using deep neural network architectures. Other models like LS-GKM [11], which evolved from the original gkm-SVM [12] model, have leveraged more conventional machine learning techniques such as support vector machines (SVMs) to predict RBP binding affinity. However, all of these models have gaps in several important areas: Firstly, in the case of the conventional machine learning models, there is typically *too* much focus on optimal k-mer sequence matching; this can entirely miss distributed recognition scenarios involving multiple nearby weaker binding motifs, and fails to account for distal factors affecting binding *in vivo*. One example was highlighted by mCross [13], which combined k-mer matching with precise crosslinking position registration and showed that SRSF1 can often recognize clusters of GGA half sites in addition to its canonical GGAGGA motif; these cases of weak motif clusters are difficult to capture with a singular k-mer focus. Secondly, when the feature set is extremely large and network architecture extremely deep or complex, as in the case of many popular neural network models, there is a frequent danger of lessened model interpretability. And thirdly, model predictions tend to be only for presence of binding within a region, with any possible downstream localization scripts being inefficient and slow to use to generate localized RBP binding maps at large scale.

Here we describe POLARIS, an integrated convolutional neural network (CNN) model that utilizes not only sequence context, but also target gene expression levels and regional transcript annotation in order to generate a more accurate model for RBP binding prediction. By incorporating these additional factors into the feature set and simplifying the network architecture, POLARIS is able to reach excellent validation set performance while remaining an interpretable and biologically grounded model. Finally, the built-in GradCAM module allows localization at single bp-resolution of the RBP signal on the original input sequence, in a way that is mathematically efficient and can be run at large scale to generate fine-tuned RBP binding maps.

## Results

### Binding model structure and predictors

POLARIS utilizes a convolutional neural network (CNN) as the engine to drive its sequence pattern recognition module. CNNs were originally designed for computer vision tasks such as handwritten digit interpretation [14–16], and more recently have been applied with success in many areas of genomics and structural biology. The principle use of convolutional layers is their capability to extract hierarchical features, or nonlinear spatial local patterns, from images or sequence data. In effect, successive layers perform new feature creation using the feature set of the previous layer’s nodes, with the final layer determining a prediction based on advanced compound features. When building a modern CNN, each convolution layer is typically immediately followed by a ReLu Activation function, which adds nonlinearity and sharpens features with positive predictive values, Max Pooling, which adds shift invariance and helps computability [17], and random Dropout, which acts to regularize the training process. Model weights and biases are all updated each epoch in the training process via backpropagation, which converts the complicated neural network update task into a tenable stochastic gradient descent problem [18].

POLARIS’ design follows this classical convolutional neural net framework: the first set of convolutional and max pooling layers serve to learn precise sequence motifs, while the second set of layers captures overarching patterns in the sequence (Figure 1). Max pooling and Dropout, as well as L2 regularization within the convolutional layers, are used to generalize training and make the model more robust. Because presence of a sequence-binding motif alone is insufficient to indicate RBP binding in tissues where the gene is not expressed - RBPs only bind to certain genes in certain tissue contexts - gene expression level is used to help guide the likelihood of actual RBP binding *in vivo*. On the other hand, inclusion of transcript regional annotation data takes advantage of the varying sequence properties at different regions of the transcript: for example, GC content of 5’UTR regions is typically higher than that of 3’UTR regions [19], which can heavily influence motif recognition and RBP binding affinity.

**Figure 1.**
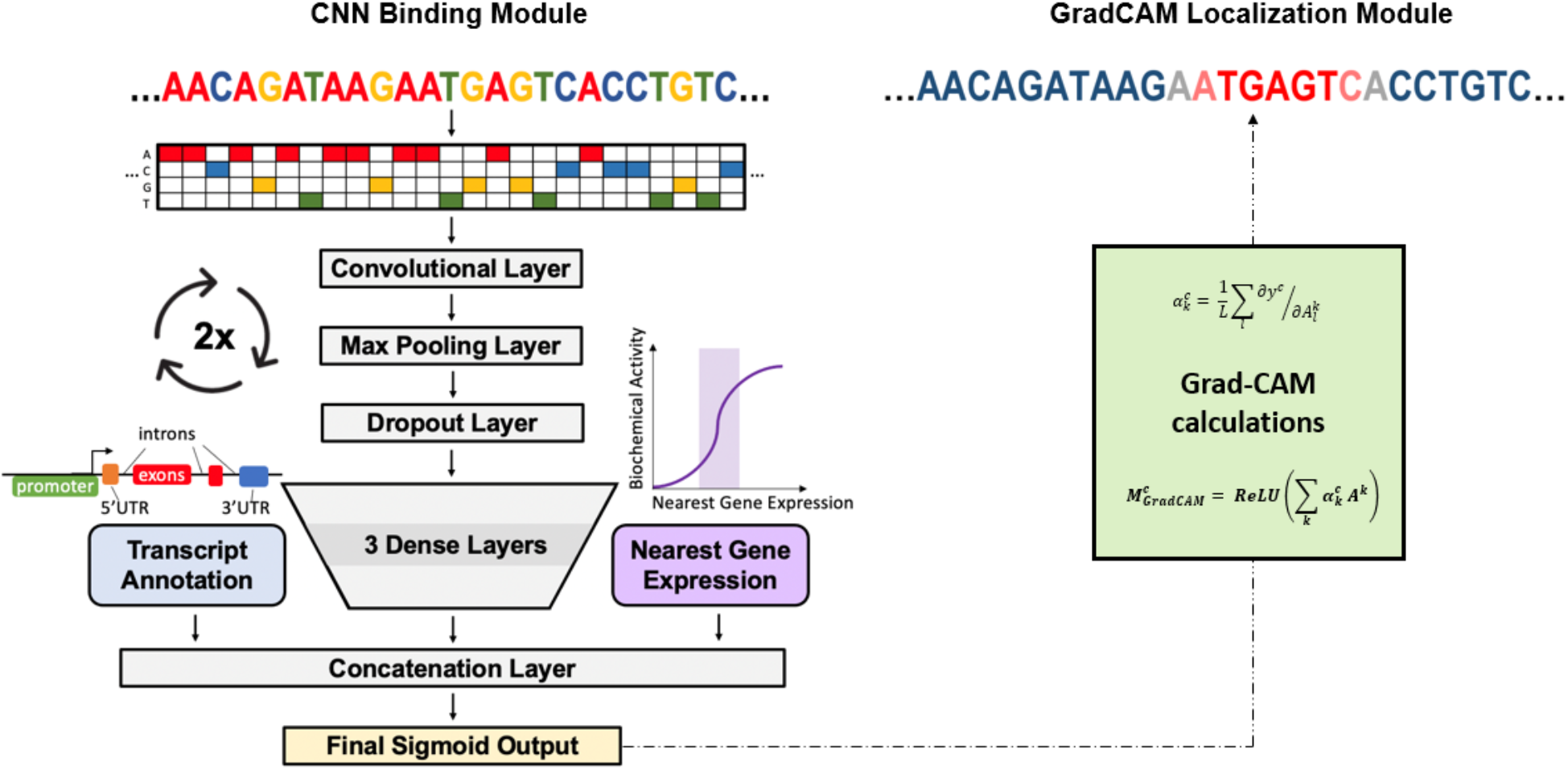
POLARIS model architecture. The binding module uses a classic two-layer convolutional neural network (CNN) structure, with max pooling and dropout layers after each convolution, and eventual integration of transcript region type annotation and nearest gene expression before the final Sigmoid binding output. The GradCAM localization module calculates the gradient of each convolutional filter with respect to this output, and maps the bp-specific contribution to the output of each position in the input sequence. In other words, it highlights the RBP binding motif(s) driving recognition of the full window as a binding region for a particular RBP.

### Model training and evaluation

POLARIS was trained using eCLIP RBP assay peaks, taken from ENCODE database.^14^ RBPs generally had ≫10k peaks each (median: 28,237 peaks, mean; 38,671 peaks, sd: 33,087 peaks), a large enough N to suggest feasibility of deep learning (Supplementary Figure S1). These peaks were further processed with CLIP Tool Kit (CTK) [20] into narrower, higher confidence sets of *in vivo* binding sites for each RBP; these *positive* training data regions indicate true RBP binding either within or close to them. *Negative* sequences for each RBP were sampled at random from transcribed regions of the genome, under the constraint that the overall GC content distribution of the negatives match that of the corresponding positives (Supplementary Figure S2).

In addition to genomic sequence inputs, POLARIS also includes input channels for target gene expression levels of the corresponding cell type where the eCLIP sequence data was generated, and transcript regional annotation type (3’UTR, 5’UTR, introns, exons, promoters, proximal 1-5kb intergenic sequence, or no annotation) of each sequence. Regional annotation type was determined by the longest transcript in the given region from GENCODE, and the gene expression data is logscale transcript parts per million (TPM) values from the RNA-seq data by ENCODE. We used all of these inputs to train POLARIS on 112 unique RBPs from the adrenal gland, HepG2, and K562 cell lines, for a total of 160 separate RBP models (if a RBP had data from multiple cell lines, they were trained and considered separately).

We randomly partitioned the eCLIP data to training (80%) and testing (20%), with balanced positives and negatives. During training, we used five-fold cross-validation to evaluate models for each RBP. The binding model with the highest mean AUC in cross-validation was selected as the optimal model for each RBP. Same-cell model AUCs referenced in this paper refer to validation set performance of these final RBP models for the most fair and generalizable metric. The POLARIS model was implemented in Tensorflow 2.2 with Keras API, with the entire training process taking around 5-10 min per RBP on an NVIDIA GTX 1080 GPU with 2560 cores and 8 GB memory.

### Performance of POLARIS in predicting RBP binding affinity

We find that POLARIS is able to successfully predict binding, with a median area under the curve (AUC) of 0.957 across all 160 RBPs analyzed (validation set, 5-fold CV, same-cell). To understand performance variance across RBPs, we plotted the distribution of binding prediction AUCs for these 160 RBPs (Figure 2a). As expected, POLARIS’ model performance improves as the sample size for an RBP increases (Figure 2b).

**Figure 2.**
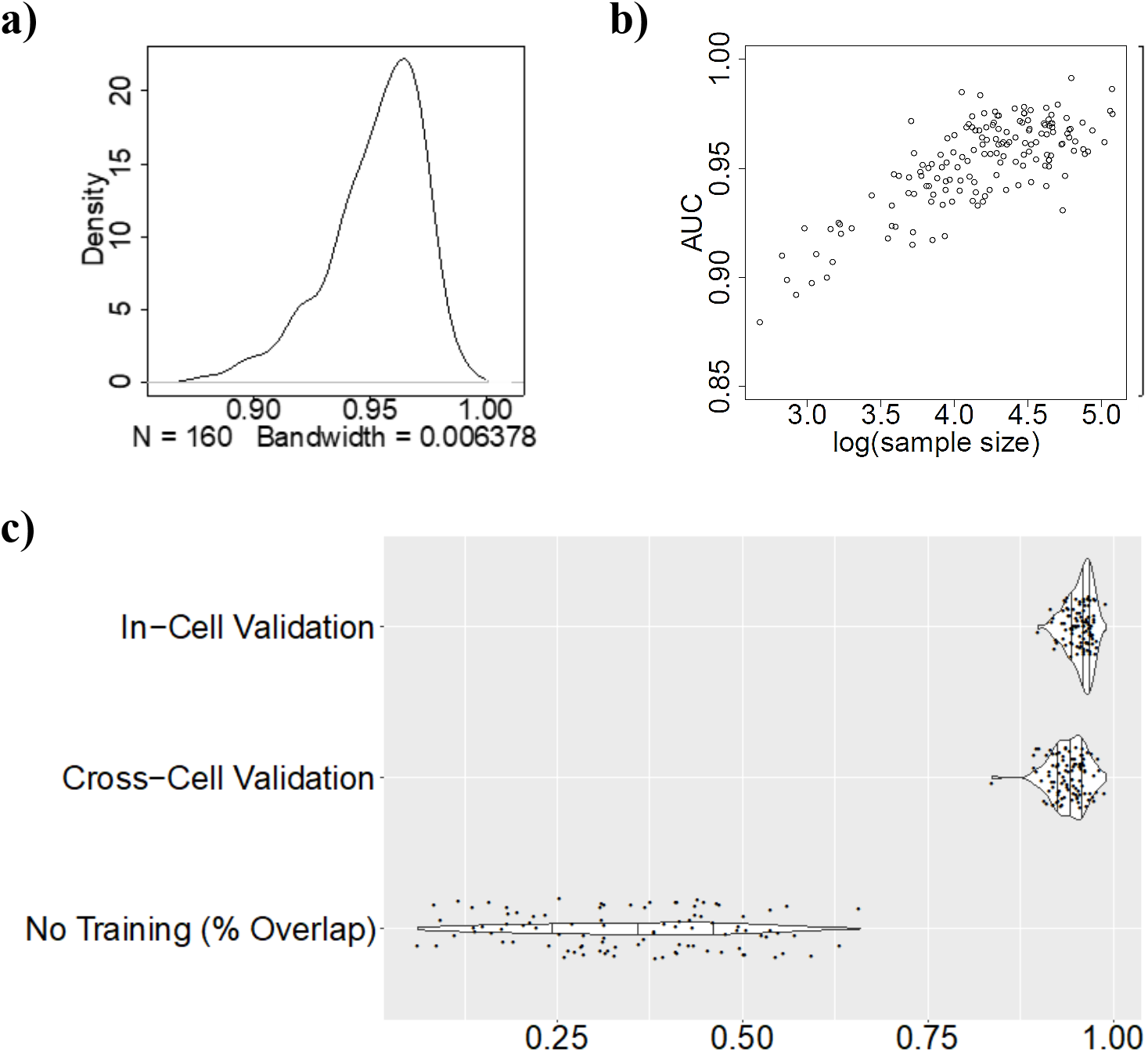
Performance of POLARIS on predicting RBP binding affinity using in-cell 5-fold cross validation. **a)** Density distribution of area under receiver operating characteristic curves (AUC) for each RBP binding prediction across all 160 RBPs. **b)** AUC of each RBP plotted against its respective sample size in log scale. **c)** Comparison of in-cell vs. cross-cell comparisons (first two rows). The final row “No Training” shows cross-cell accuracy with a naïve baseline model (negative control) that predicts K562 binding status exactly from HepG2 (and vice versa).

A major goal of POLARIS is to predict cell-type specific RBP binding. To evaluate this, we tested the trained models for each given RBP across cell lines. For example, we trained a QKI binding model on data from the HepG2 cell line, then tested the model using data from the K562 cell line, and vice versa. POLARIS was still able to achieve a high AUC of 0.942 using cross-cell method of validation (Figure 2c, first two rows). As a baseline method, we used the eCLIP sites from one cell line (HepG2, K562) to directly predict binding in the other - without any actual neural network training - to see how much inherent overlap there was in the binding peaks. The final row of Figure 2c shows that this baseline method performs poorly, with a median AUC of only 0.778.

### Evaluation of prediction feature importance

When designing the model framework for POLARIS, we saw that including features based on prior knowledge of biological mechanisms, such as target gene expression and transcript regional annotation categories, boosted model performance (Figure 3a). Addition of the regional annotation feature to the base CNN model increased median AUC from 0.899 to 0.932. Alternatively, addition of gene expression values increased median AUC from 0.899 to 0.945. The best validation set performance was achieved with inclusion of both features (median AUC = 0.957), which is the setup used in the final POLARIS model.

**Figure 3.**
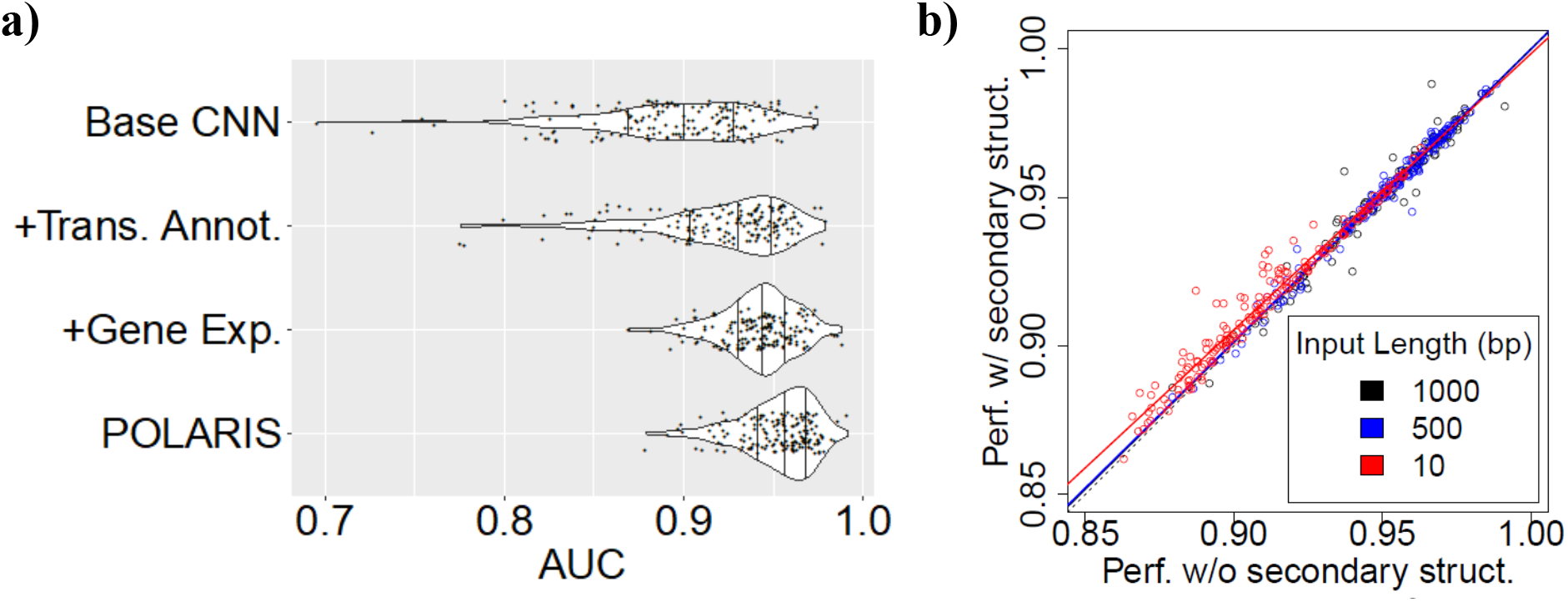
Evaluation of POLARIS model features and secondary structure impacts. **a)** RBP binding prediction improves when explicitly adding gene expression and transcript annotation information as inputs. The final binding prediction model used in SUPRNOVA incorporates both gene expression and annotation inputs. **b)** Performance (measured as AUC) when including secondary structure context (hairpin loop, internal loop, etc.) is slightly improved when the input sequence length is small. This is indicated by a shallower slope than 1 for the red 10bp line. However, once the input sequence length is increased, included secondary structure information no longer affects POLARIS’s performance.

### Effects of adding predicted secondary structure of mRNA

RNA secondary structure is a major factor of binding affinity of certain RBPs [21]. We followed a technique presented in Koo et al. [22] to investigate if including secondary structure information as input can improve binding prediction. After using RNAplfold [23] (with Kazan, H.’s modified script [24]) to annotate sequences with predicted secondary structure information, we fed this information into our model as an additional input channel (similar to the channels for gene expression and regional annotation). Simpler, motif-only models such as PWMs, k-mer SVMs, and small-window neural networks see a boost in prediction performance with addition of secondary structure feature [22, 24, 25]. Indeed, even POLARIS saw a performance boost with RNAplfold when limited to a more narrow 10bp window size. However, inclusion of these structure profiles did not boost the performance of the POLARIS models with sufficiently large input sequence length (Figure 3b).

### Comparison with other methods

Besides a standard CNN, a recurrent neural network (RNN) hybrid model was also considered for the core POLARIS framework. Based on the work of Quang et al. [10], which highlighted the potential benefits of long short-term memory (LSTM) layers, we tested a CNN-LSTM hybrid version of POLARIS that replaced the second convolutional layer with an LSTM layer. The input data was exactly the same as what we had used to evaluate the CNN architecture; however, despite taking longer to train, this hybrid model did not provide any performance gain, only achieving median AUC of 0.933.

POLARIS outperforms previously published methods such as the gapped k-mer based SVM model, LS-GKM (from Lee, D. et al. [11], based on the original gkmSVM [12]), and the standard CNN-based DeepSEA model from Olga Troyanskaya’s group [8] that Seqweaver utilizes for its post-transcriptional modelling. After re-training these methods from scratch using the same input data as POLARIS and performing 5-fold cross validation, Seqweaver achieves a median AUC of 0.871, and LS-GKM reaches a median AUC of 0.892 (Figure 4a). These are more comparable to the performance of our basic model framework (median AUC = 0.900), which did not include expression or annotation data. We also tested a version of Seqweaver into which we integrated our exact expression and region type tensors used in POLARIS (labeled “Seqweaver+” in Figure 4a). This brought results significantly closer (median AUC = 0.931), but overall POLARIS was still more consistent and had higher average performance (median AUC=0.957). We show RBP-specific results for three examples: RBFOX2, ILF3, and QKI, with each trained in HepG2 and tested in K562 (Figure 4b,c,d). In ILF3 and QKI in particular, each with N ~ 30k eCLIP peaks, we see some of the most significant performance improvements over the competing models.

**Figure 4.**
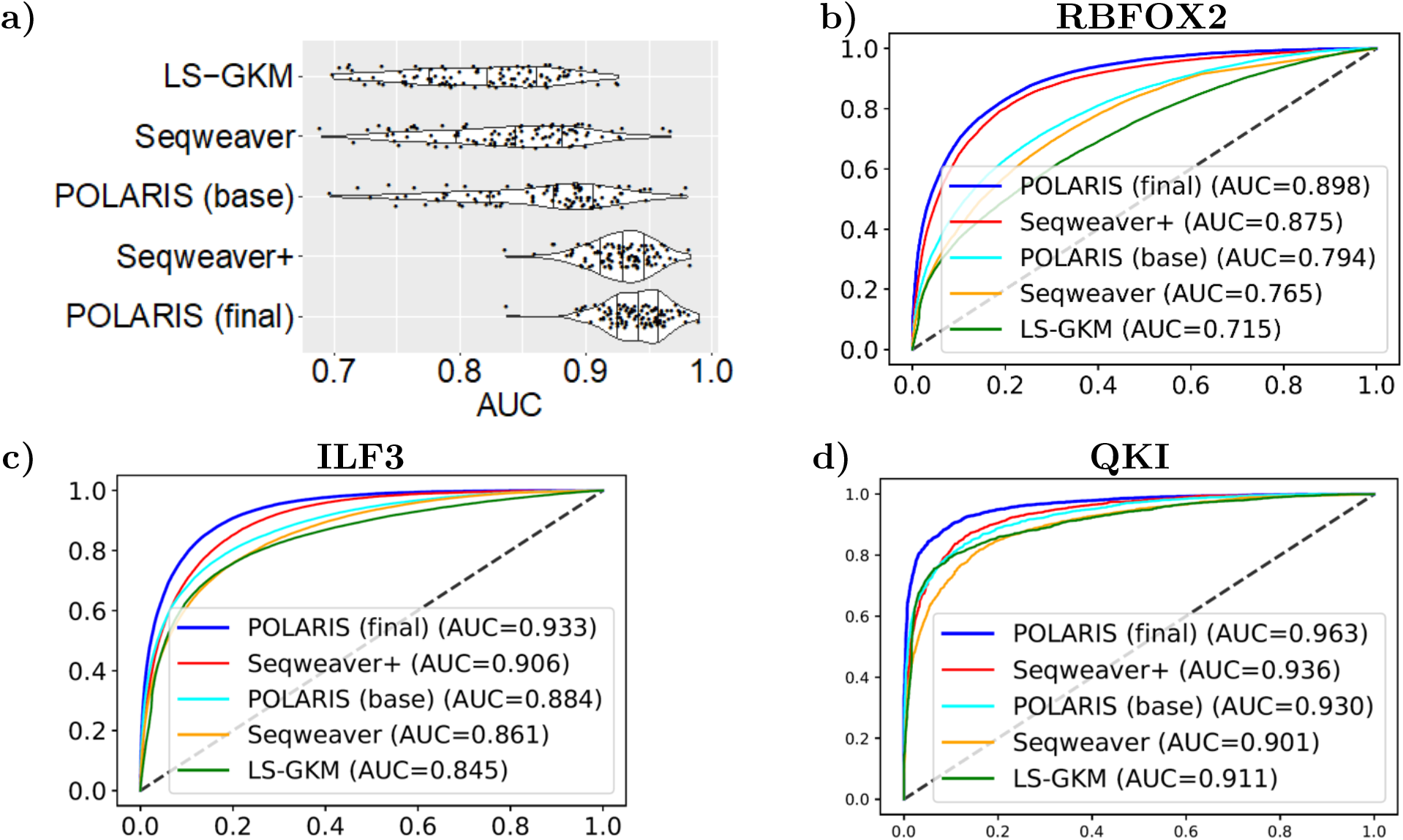
Performance of POLARIS compared to other computational methods for predicting RBP binding affinity. **a)** Violin plot showing distribution comparison of all methods’ cross-cell performance results. Depending on the RBP, cross-cell performance of Seqweaver + expression + region type annotation got near POLARIS, but overall POLARIS was more consistent and had higher average performance. **b,c,d)** Specific examples for RBFOX2, ILF3, and QKI, respectively: all were trained in HepG2 and tested in K562. In ILF3 and QKI in particular, each with N ~ 30k eCLIP peaks, we see some of the best performance improvements over the competing models.

### GradCAM results for determining localized affinity and motif discovery

Finally, we used a one-dimensional version of gradient-weighted class activation mapping (GradCAM) [26] to localize sites within an original input sequence that are informative for binding prediction (Figure 1). The powerful algorithm can robustly highlight causal subsections of input (roughly corresponding to RBP binding motifs in our case), regardless of how many layers the model contains or how the neural network architecture is set up. It also allows our localization method to be mathematically efficient, making it possible to run at large scale to generate fine-tuned binding maps for all available RBPs at once.

We implemented the GradCAM algorithm as a single reverse pass once the binding module determines an overall prediction for the full window (see Methods for full details). Recovery of canonical motifs, for two representative examples (RBPs RBFOX2 and EFTUD2), are shown in Figures 5a,b. We qualitatively compared the highest activating regions for each RBP’s sequences to their respective known canonical motifs (based on mCrossBase[13]), and found a high recovery of known motifs, as well as plausible novel candidate motifs for RBPs with no available motif data.

**Figure 5.**
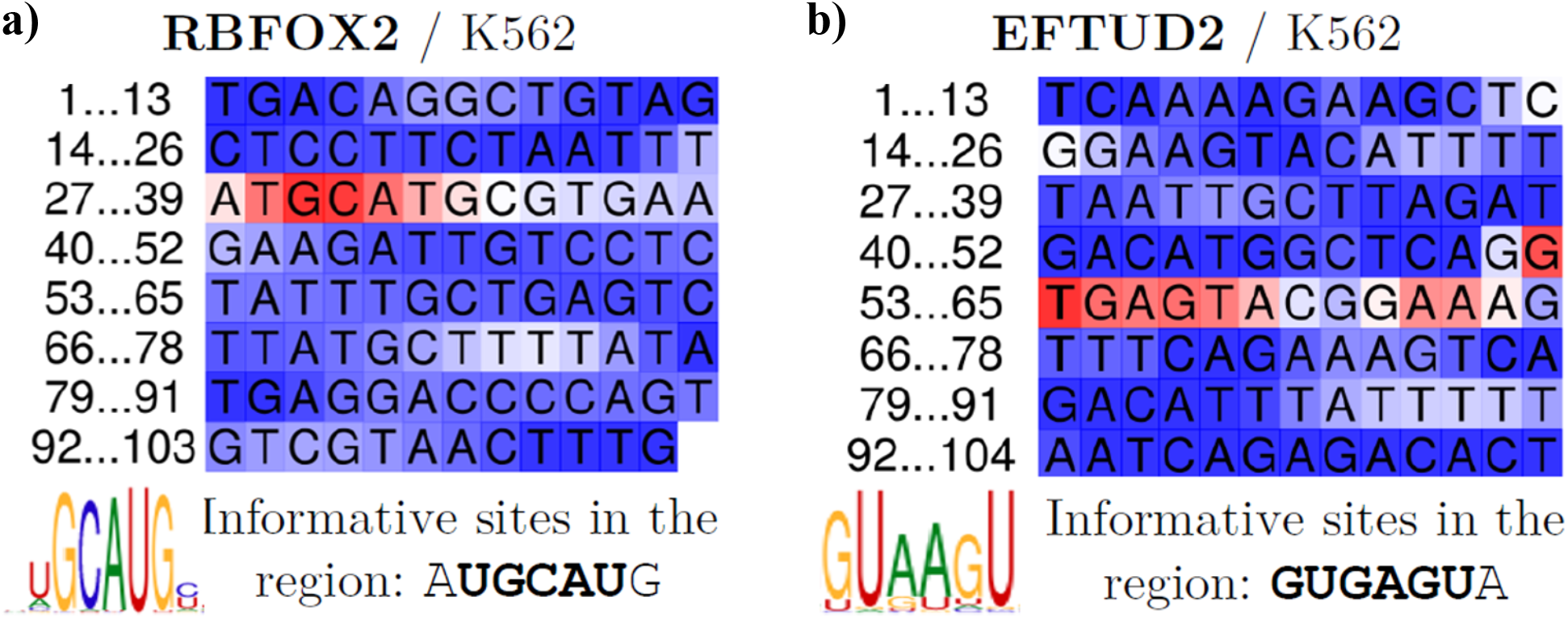
Recovery of canonical motifs using POLARIS’ GradCAM module. Shown are heatmaps for two representative RBPs: RBFOX2 **(a)** and EFTUD2 **(b)**; sequence is shown in multiple rows for visibility, but are 1-dimensional character strings. Canonical binding sites and sequence logos have been taken from mCrossBase.

### Investigating strong POLARIS performance for weak motifs with GradCAM

It is natural that POLARIS performs well when trained on RBPs whose binding motifs are especially consistent and clear to separate from surrounding genomic sequence. Thus, a naive expectation is that validation AUC positively correlates with information content of sequence logos, a proxy of motif quality. The information content for each RBP was calculated by considering the top ten strongest motif versions most associated with the given RBP; we obtained these from mCrossBase database, which makes accessible results from the aforementioned mCross paper [13]. In order to obtain a single information content metric for each RBP, we selected the maximum total information content value (summed across positions) out of all of the candidate motifs.

However, when we made a scatterplot of this relationship (Figure 6a), we found an observation of POLARIS’ performance that was not trivial to explain: although the general positive correlation trend was confirmed, there was a group of RBPs in the top-left corner with low motif information content that nevertheless had very strong performance (AUC threshold >0.95 and AUC/IC ratio of .09; highlighted in red).

**Figure 6.**
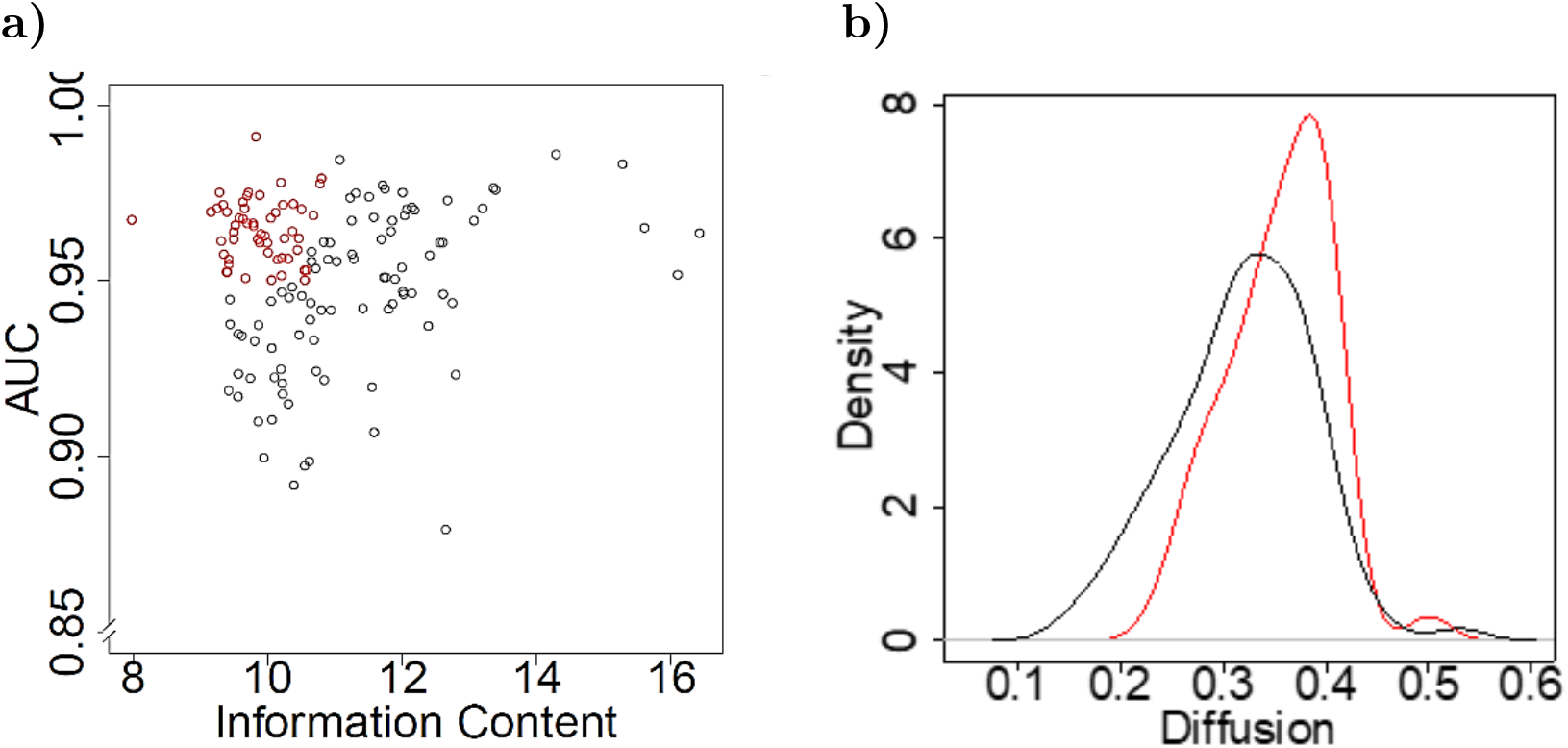
Investigating strong POLARIS performance for weak motifs. **a)** AUC of each RBP plotted against the information content for each given RBP’s binding canonical binding motif, taken from mCrossBase. Highlighted in red are the weak motifs that still have very strong performance (AUC threshold >0.95 and AUC/IC ratio of .09) **b)** Density of diffusion/entropy for red RBP models vs. black ones, using the same color scheme as panel (a): weaker motifs generally do have more spread of binding score in the window, indicating multiple binding sites in the region.

Our working hypothesis to explain this was that RBPs with weaker motifs could have multiple binding sites per binding window, to raise the total regional information content and enable post-transcriptional regulatory control with the same level of precision as RBPs with strong motifs. It was previously shown that this could be the case in the mCross paper [13], which found that the RBP SRSF1 can recognize clusters of GGA half-sites in addition to its canonical GGAGGA motif. We investigated the over-performing (red) RBP models by considering density of diffusion/entropy for them vs. the other (black) ones; calculations for this step are detailed in Methods, Equation 3. We found that weaker motifs generally *did* have more spread of binding score in the window, with a clear enrichment of their RBPs at higher diffusion values (Figure 6b). This indicates multiple binding sites in the region, and confirms the plausibility of our hypothesis.

## Discussion

In this study, we presented POLARIS (**P**rediction **O**f **L**ocalized **A**ffinity for **R**BPs **I**n **S**equence), a new deep-learning method that is an integration of a CNN to predict RBP binding to the transcribed genome from genomic sequence, nearest gene expressions, and region type annotations, with a GradCAM implementation for efficient localization of this signal backpropagation to individual sequence positions. We demonstrated that the model is able to achieve very strong RBP binding prediction performance (~0.957 median AUC) that outperforms traditional sequence-only based prediction models and competing neural networks. Indeed, the results of DeepSEA (median AUC = 0.871) and LS-GKM (median AUC = 0.892) on our held-out validation set were more comparable to the performance of our basic POLARIS model framework, which does not include expression or region annotation data; this suggests that the biological mechanisms that drive feature selection in POLARIS are a big part of what helps our model outperform existing methods. By incorporating these additional factors (gene expression, region type) into its base feature set, POLARIS is able to reach optimal validation set performance with fewer convolutional filters, and is consequently easier to parse and understand. Thus, despite utilizing neural networks, POLARIS is both interpretable and biologically grounded, important qualities to be able to verify results, investigate the genetic underpinnings driving model predictions, and make educated application of the model in downstream analyses.

Regarding model architecture, POLARIS model layers have clear roles: upper convolutional layers handle the bulk of RBP motif learning, later convolutional layers are responsible for incorporating more distal effects like global RNA secondary structure, and the intermediate dropout and max pooling layers help select representative data, minimize risk of overfitting, and enhance learning efficiency. Keeping POLARIS extremely condensed was not a top priority, since even the full 1kb models do not take up much storage space or runtime on modern machines or clusters: thus, we decided to learn a large number of informative filters (100) for each RBP individually, rather than dilute model power with individual filters for each RBP within one structure. At the same time, we use various regularization techniques throughout the model (L2 regularization in convolutional layers, followed by Max Pooling and Dropout), which combined with our relatively simple architecture help reduce overfitting risk despite a large number of total trainable parameters. Our experiments with a recurrent neural network (RNN) hybrid model did not provide any notable performance gain (median AUC only 0.933). This implies that our final CNN-based POLARIS model is sufficient in capturing hidden sequential information that is actually useful for *in vivo* binding prediction.

We took several important steps to optimize training and guide the model towards learning real biological and genetic properties. For instance, without the step of accounting for GC content in the randomly sampled negative training sequences, the prediction model could inadvertently learn confounding patterns in the data that are not directly relevant to RBP binding. We also minimized overfitting risk within our robust training and evaluation procedures, although we caution that all of the original RBP binding data was generated exclusively with eCLIP experiments from ENCODE – thus, it is not possible to rule out that some aspect of POLARIS models artifacts from eCLIP experimental protocol. We contend that this is unlikely to be significant, and future testing on alternative data sources will confirm the full generalizability of our model to arbitrary sources of sequence binding data. Finally, our reported POLARIS AUCs were taken from validation set performance, which was entirely isolated during model training and selection. Performance remained high (median AUC ~ 0.942) even when using cross-cell evaluation, despite eCLIP overlap being relatively low and therefore making this a challenging test. Altogether, these results indicate that our model successfully learned sequence motifs and general in vivo binding affinity patterns for each given RBP, rather than over-fitted patterns unique to a certain tissue or region context.

In accordance with expectations, we also saw that as the sample size for an RBP increases, so too does its model performance: this is because neural networks require sufficiently large training sets to be able to accurately learn patterns within the dataset, so with more data, the model has more information to learn from to outperform competing models at prediction of in vivo RBP binding. In this way, use-cases involving simultaneous integration of multiple RBPs will have more stable and consistent behavior across all analyzed RBPs. POLARIS model performance being not only high but also robust across RBPs and cell line allows for extrapolation and utility in tissues other than those necessarily present in the training set: this directly counters the critical hurdle of sparse data, which inevitably presents itself when trying to build such models with the goal of generalizability.

We show that our performance gain over competing models is plausible in two key ways: 1. evidence that the performance gain over simpler competing models such as PWMs and SVMs can be attributed to implicit learning of real biological factors such as RNA secondary structure, and 2. Direct extraction and visualization of informative RBP binding motifs, using our implementation of gradient-weighted class activation mapping. Although POLARIS saw a performance boost with RNAplfold when limited to a narrow 10bp window size, inclusion of these structure profiles did not boost 1kb POLARIS model performance when given sufficiently large input sequence length. As suggested by Koo, et al., this indicates that the original model is already able to implicitly capture this information within later convolutional filters. While we caution that the dynamic nature of secondary structure and inaccuracy of prediction can confound the analysis, this apparent secondary structure learning from distal sequence represents a very important advantage when modeling *in vivo* RBP binding instead of merely motif match/mismatch. Another helpful factor could be our more careful curation of representative training data, including upstream fine-tuning of eCLIP peaks with CLIP Tool Kit (to improve resolution of protein-RNA interactions by determination of exact crosslink sites and connection of peak valleys) and RBP-specific training with custom GC-balanced negative sets. These steps help the model focus on learning only representative motifs and relevant distal features for binding prediction, rather than noise or biases in the data. It is common in the field to use random eCLIP sites of other RBPs as negatives for a given RBP, in order to control for eCLIP experimental protocol and regional preferences. However, we believe that our transcribed, GC-balanced negative regions could actually account *better* for the most prominent of these factors for each individually trained RBP, while simultaneously enabling our model to minimize any biases against RBP complexes, the primary way in which RBPs physically act upon transcripts in reality. Additional validations on alternate sources of RBP sequence binding data, such as older cross-linking immunoprecipitation (CLIP-seq) data, will be required to verify this intuition.

Finally, POLARIS’ GradCAM module provides a method to interpret the prediction in a way similar to conventional binding motifs without the limitation of linearity. This ability to efficiently backpropogate binding signal to subsections of the input sequence driving the prediction (at single base pair resolution) is a novel addition to the field of genomic binding modeling with CNNs. It is useful both for localizing overall RBP binding predictions and for quickly extracting biologically-grounded RBP binding motifs, equally canonical and novel ones. Overall, with its focus on localized prediction, POLARIS is especially well suited for in vivo binding motif interpretability and downstream variant effect prediction. In particular, we expect the POLARIS model can be a component of a method for predicting functional impact of noncoding variants useful for genetic studies.

## Methods

### Data processing for RBP binding model

We obtained eCLIP (enhanced crosslinking and immunoprecipitation) RBP binding data from ENCODE [5]. eCLIP data is stored as rows of peaks, specifying chromosome, start, and end position of each peak together with an associated p-value. In total, we acquired binding data for 112 unique RBPs from the adrenal gland, HepG2, and K562 cell lines, for a total of 160 separate RBP eCLIP peak files. Our final positive dataset for the RBP binding model was generated from this eCLIP data by first removing singular peaks that are not within 500bp of any other peak for a given RBP, and also excluding peaks with p-values > .01 to minimize the presence of noisy data. These peaks were further processed with CLIP Tool Kit (CTK) [20] into narrower, higher confidence sets of *in vivo* binding sites for each RBP; these *positive* training data regions indicate true RBP binding either within or close to them. Finally, the regions were padded (equally) on both sides to 1kb total sequence length each in order to ensure that the binding motifs were contained within the input contexts, and to be able to learn binding effects from more distal sequence elements.

The negative dataset was generated by randomly sampling transcribed sequences (pre-mRNA transcripts) along the hg19 refGene genome from UCSC. The sequences are also constrained to the full peak ranges of their corresponding cell line, and only include at most 500bp of non-transcribed regions. Since the positive sequences were padded up to 1kb, the negative sequences were allowed to overlap with positive sequences, by at most 200bp. These random sequences were also evenly distributed between the positive and negative strands, which matches the strand distribution of the positive dataset.

Each one of these positive and negative sequences was then one-hot encoded and combined into a 3D tensor, thereby creating a vectorized representation of the sequence strings. Since the sequences have variable length, empty space was added to the end of shorter sequences as padding in order to ensure consistent dimensionality throughout the tensor.

### RBP binding model architecture details

In POLARIS’ CNN, the convolution layers compute output by one-dimensional convolution operation with a specified kernel size and number of filters. Then, the pooling layers compute the maximum value in a specified window of spatially adjacent convolution layer outputs for each kernel. The fully connected dense layers on top of the second convolution receives input from all of the outputs from the previous layer, thereby integrating information from the full sequence length. The dense layers perform a rectified linear (ReLU) activation to the hidden unit cells:

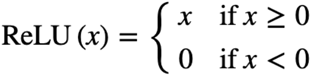

Finally, the sigmoid output layer makes a prediction of whether or not the RBP will bind to a given sequence, and scales the prediction to the 0-1 range by the sigmoid function:

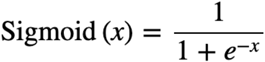

In order to prevent overfitting, a proportion of outputs were randomly set to zero at some of the layers. A dropout proportion of 15% was added after layers 2, 4, and 7. Both of the convolution layers also had an L2 regularization term of .01, which again, was used to minimize overfitting and add some robustness to the training process. Finally, this model uses a stochastic gradient descent optimizer and a binary cross-entropy objective loss function:

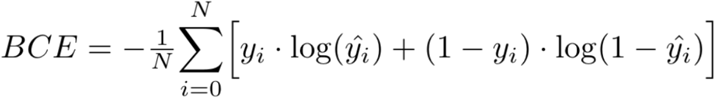

This model was trained with a batch size of 16 and 10 epochs, which was empirically observed to lead to a plateau in performance. Specific architecture and final hyperparameters are outlined on our GitHub page (see Appendix).

Both of the RNN models use a binary cross entropy loss function and an RMSprop optimizer. They were trained with a batch size of 100 and ran for 10 epochs. Specific architecture and hyperparameters are again outlined on our GitHub page (see Appendix).

In the current version of POLARIS, individual RBP models do not share layer parameters amongst themselves; we made this design decision to concentrate power to detect each individual RBP, rather than force the filters to multi-task (with less controlled negative set data, since GC balancing is unique to each RBP). However, after individual training these models can trivially be joined together with a shared input layer, to have a more condensed full model that simultaneously predicts binding for all RBPs at once.

All neural network models mentioned above were built using the Python keras package with a GPU-accelerated Tensorflow (v2.2) backend, and trained using a machine with one Nvidia GTX1080 graphics card (2560 CUDA cores) and 8 GB RAM. With these specifications, the full POLARIS training process takes between 5-10 min per RBP.

### Annotations and gene sets

Variants were generally annotated using ANNOVAR (v2017-07-17). Region type annotation was done with the annotatr R package, from the Bioconductor suite.

### Gradient-weighted Class Activation Mapping (GradCAM) implementation

Class activation mapping (and GradCAM in particular) was originally developed for image recognition tasks; however, like CNNs themselves, the algorithms translate very well to the case of binding motif modeling on 1-dimensional sequence. The method produces a coarse localization map highlighting the important regions in the input image/sequence for predicting the concept class (specific RBP binding, in our case). Given an input, GradCAM captures the outputted feature map of the convolution layer and weights every filter in that feature map by the gradient of the class with respect to the filter. Intuitively, filter activation is balanced by the importance of each filter to the output class, and this weighted sum creates a spatial map of class activation by the input: we can then interpolate/stretch the resulting heatmap to original input size and overlay onto the image/sequence. In other words, GradCAM allows us to robustly highlight the causal subsections of input regardless of how many layers the model contains or how the neural network architecture is set up. It also allows our binding localization implementation to be mathematically efficient, requiring only the one original pass through the network per input; it can thus be run at large scale to generate fine-tuned binding maps for all available RBPs at once.

We implemented the Gradient-weighted Class Activation Mapping (GradCAM) algorithm design described in Selvaraju et al., 2017 [26], with the simple modification that instead of projecting onto a u x v dimensional image, we find the activation map onto a one-dimensional vector of length L base pairs for the case of our RBP binding model (which only performs a one-dimensional sequence convolution, rather than 2D “image” convolution). In order to obtain the class-discriminative localization map 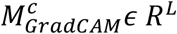 of length L base pairs for any RBP class c, we first compute the gradients via backpropagation of class c’s score ***y^c^***, with respect to feature map / filter activations ***A^k^***, i.e. ***∂y^c^/∂A^k^***. These gradients flowing back are global-average-pooled over the length dimension (indexed by l) to obtain the neuron importance weights 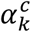:

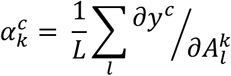

**Equation 1. Averaging class-specific filter activations to find GradCAM neuron importance**

During computation of 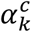 while backpropagating gradients with respect to activations, the exact computation amounts to successive matrix products of the weight matrices and the gradient with respect to activation functions till the final convolution layer that the gradients are being propagated to. Hence, this weight 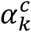 represents a partial linearization of the deep network downstream from A, and captures the “importance” of filter k for a target class c.

We perform a weighted combination of forward activation maps, and follow it by a ReLU to obtain

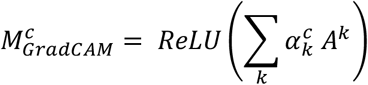

**Equation 2. Calculation of GradCAM localization by weighting forward activation maps**

Notice that this results in a coarse heatmap of the same dimensionality as the convolutional filters, but is easily stretched back up to the original input size for projection (151 positions, in the case of our RBP binding models). A ReLU is applied to the linear combination of maps because we are interested in the features that have a positive influence on the class of interest; as Selvaraju, et al. note, negative “pixels” or positions are likely to belong to other categories and GradCAM localization performance is decreased without this ReLU operation. In total, for each sequence we can end up with 160 L-length GradCAM heatmaps, one for every RBP class c, that highlight the most relevant regions of the input for placement in each respective class.

### Calculating average GradCAM Diffusion for RBPs

Diffusion for each RBP was calculated by averaging the Shannon’s entropy H(X_*q*_) of the GradCAM heatmap output for each of its known positive eCLIP binding sequences *q*, with each sequence’s entropy first normalized by total binding strength of the sequence window (sum of positional GradCAM heatmap vector X_*q*_). This normalization step was included in order to focus only on capturing RBP motif diffusion, rather than any proxy of binding motif quality (Equation 3).

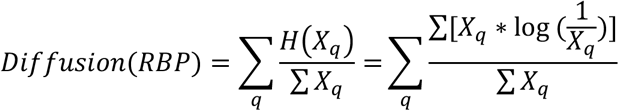

**Equation 3. Calculation of RBP by normalization of sequence Entropies by total binding strength**

## Data and Code availability

All training and test data and code are available on GitHub: https://github.com/ShenLab/noncoding

## Acknowledgements

We thank Dr. Chaolin Zhang for sharing processed eCLIP peaks and helpful discussions. The work was supported by NIH grants R01GM120609 (Y.S.) and R03HL147197 (A.K. and Y.S.).

## Author contributions

All authors contributed to data analysis, interpretation, and manuscript writing. Y.S. conceived and designed the study.

## Appendix Generating GC-balanced negative training data

POLARIS was trained using eCLIP RBP assay peaks, taken from ENCODE database.14 RBPs generally had ≫10k peaks each (median: 28,237 peaks, mean; 38,671 peaks, sd: 33,087 peaks), a large enough N to suggest feasibility of deep learning (Supplementary Figure S1).

**Supplementary Figure S1.**
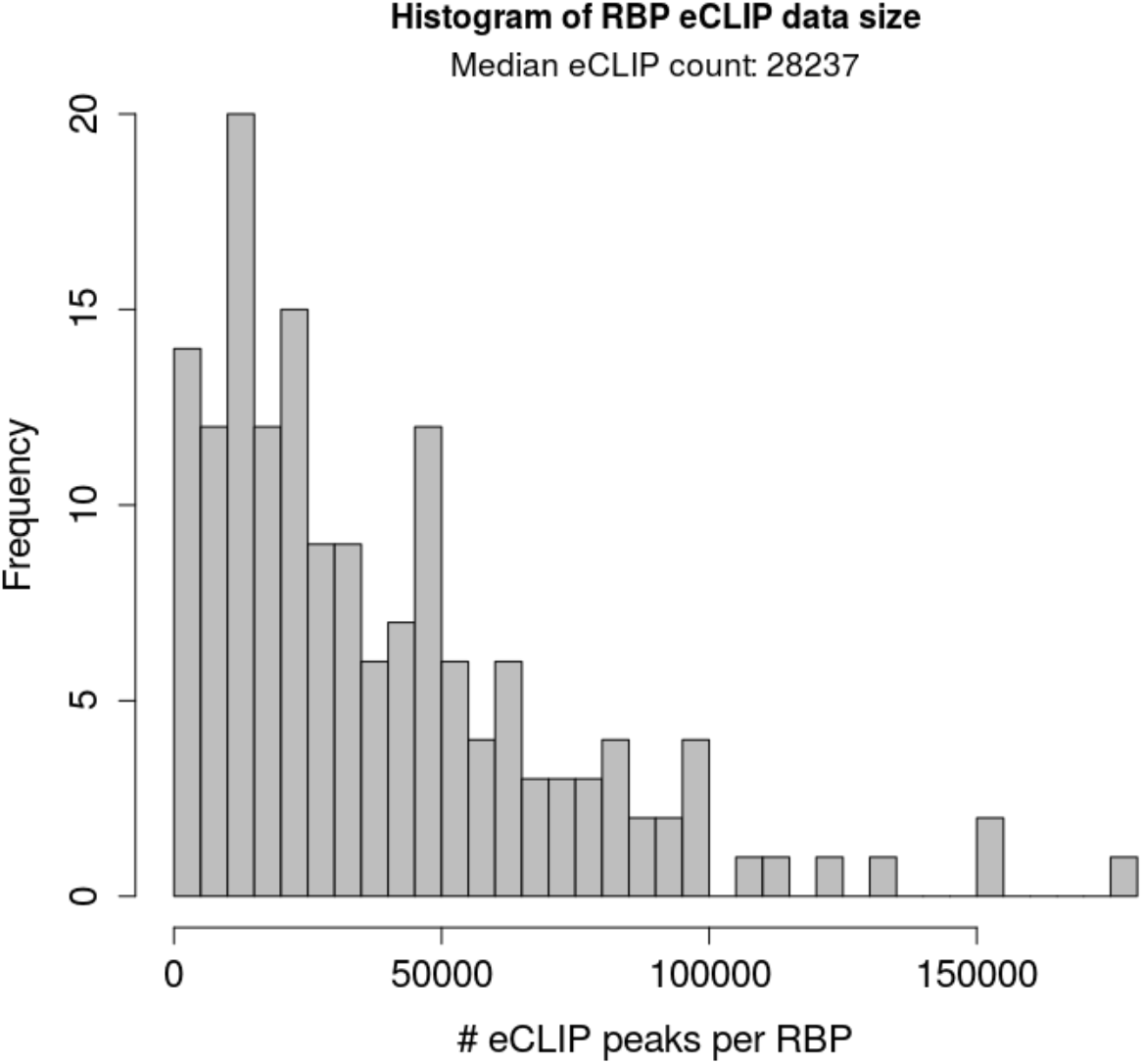
Histogram of eCLIP peak count per RBP. The majority of RBPs had N ≫ 10k peaks, indicating plausibility of applying deep learning. Median: 28,237 peaks, mean; 38,671 peaks, sd: 33,087 peaks.

These peaks were further processed with CLIP Tool Kit (CTK) [20] into narrower, higher confidence sets of *in vivo* binding sites for each RBP; these *positive* training data regions indicate true RBP binding either within or close to them. Finally, the regions were padded (equally) on both sides to 1kb total) sequence length, to be able to learn binding effects from reasonably distal sequence elements. *Negative* sequences for each P were sampled at random from transcribed regions of the genome, under the constraint that the overall GC content distribution of the negatives match that of the corresponding positives (Supplementary Figure S2).

**Supplementary Figure S2.**
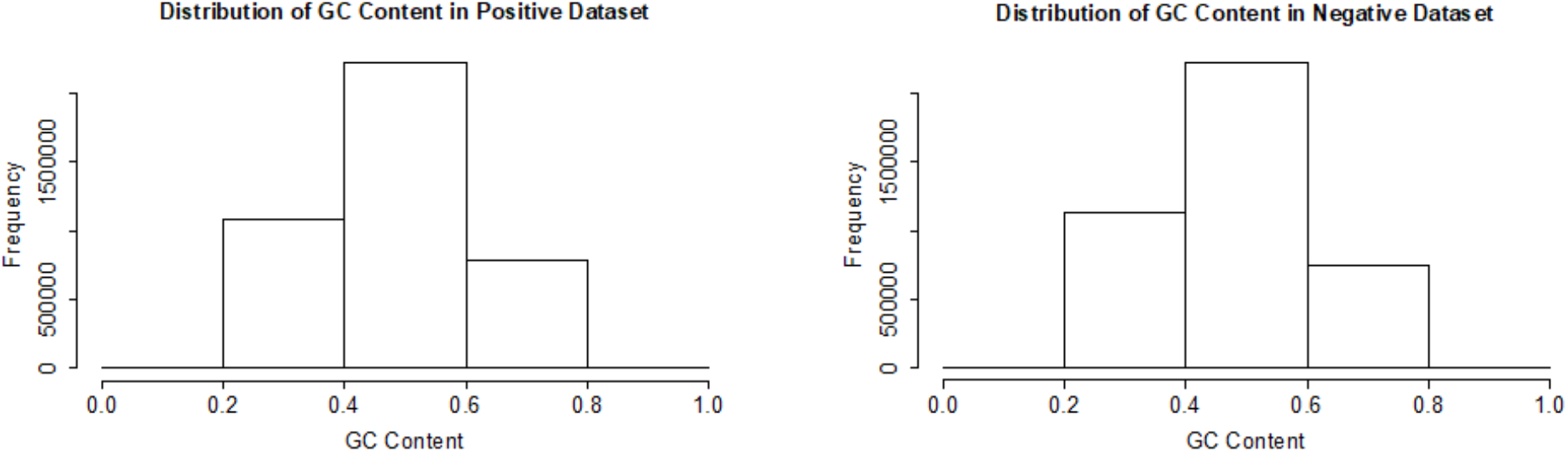
Comparison of GC content distributions. Regional GC content distributions are shown for both the positive (eCLIP) dataset **(a)**, and the sampled negative (random transcribed regions) dataset **(b)**. GC distribution is intentionally matched in the training data (for each RBP binding model), to not accidently train this feature.

## Clustering GradCAM heatmaps to binding motifs and valleys

We developed a weighted hierarchical clustering algorithm to automatically identify motifs and motif valleys from GradCAM heatmaps. Notably, our approach does not rely on fixed hyperparameter constraints on optimal motif length k or the number of motifs within a window, which can both strongly differ across RBPs and binding regions. We first create a pairwise distance matrix D that is the weighted sum of two distinct distance matrices: a) Score distance, where score is the GradCAM projection score ϵ (0,1) at every position, and b) Sequence distance, which is simply the base pair gap between every pair of positions. The relative impact of each of these matrices on D is controlled by a weight parameter α, helping guide the function to select motifs of reasonable length (higher α prioritizes score similarity over sequence proximity, encouraging consideration of longer motifs). An example distance matrix calculation for heatmap vector [1,1,1,0,0,0,1,1,1] and balanced α = 0.5 is shown below (Supplementary Equation 1).

**Supp. Equation S1.**
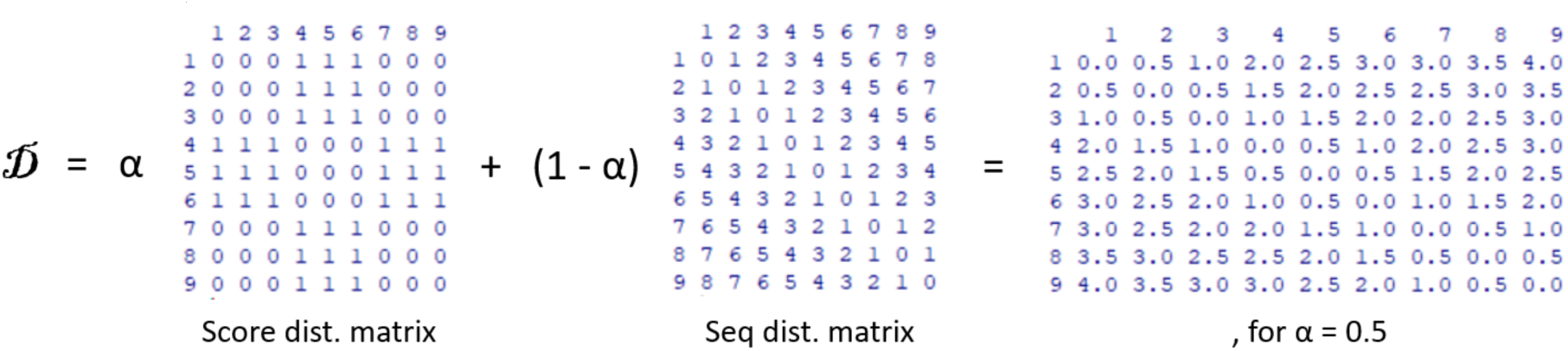
Example calculation for pairwise distance matrix D for GradCAM heatmap.

We then perform hierarchical clustering on D, producing a hierarchical tree; we found the results were most consistent when complete / furthest neighbor clustering was used, but the provided algorithm function supports average distance / UPGMA (unweighted pair group method with arithmetic mean) or any other distance aggregation function as well. We cut the tree at a fixed height h to obtain sequence clusters C. Finally, we find aggregate cluster scores S for each c ϵ C by averaging member scores within each cluster, and designate the clusters where S > m as RBP binding motifs; the remaining clusters represent connected valleys between motif plateaus.

An h of 1 represents a furthest score distance of ~1 between directly neighboring clusters for the case of α = 0.5, and worked well in practice for a wide range of α up to 0.9. We also recommend binding threshold m > 0.5, which can be increased further to prioritize specificity over sensitivity in motif discovery. Using this hyperparameter selection, we ran our algorithm on eCLIP-positive validation set regions. Qualitatively, although resolving borderline cases with low POLARIS binding prediction scores was difficult, when binding prediction score was high (generally >0.5), the clustering worked very well to extract motifs; we show an example for the important splicing regulator QKI below (Supplementary Figure S3).

**Supplementary Figure S3.**
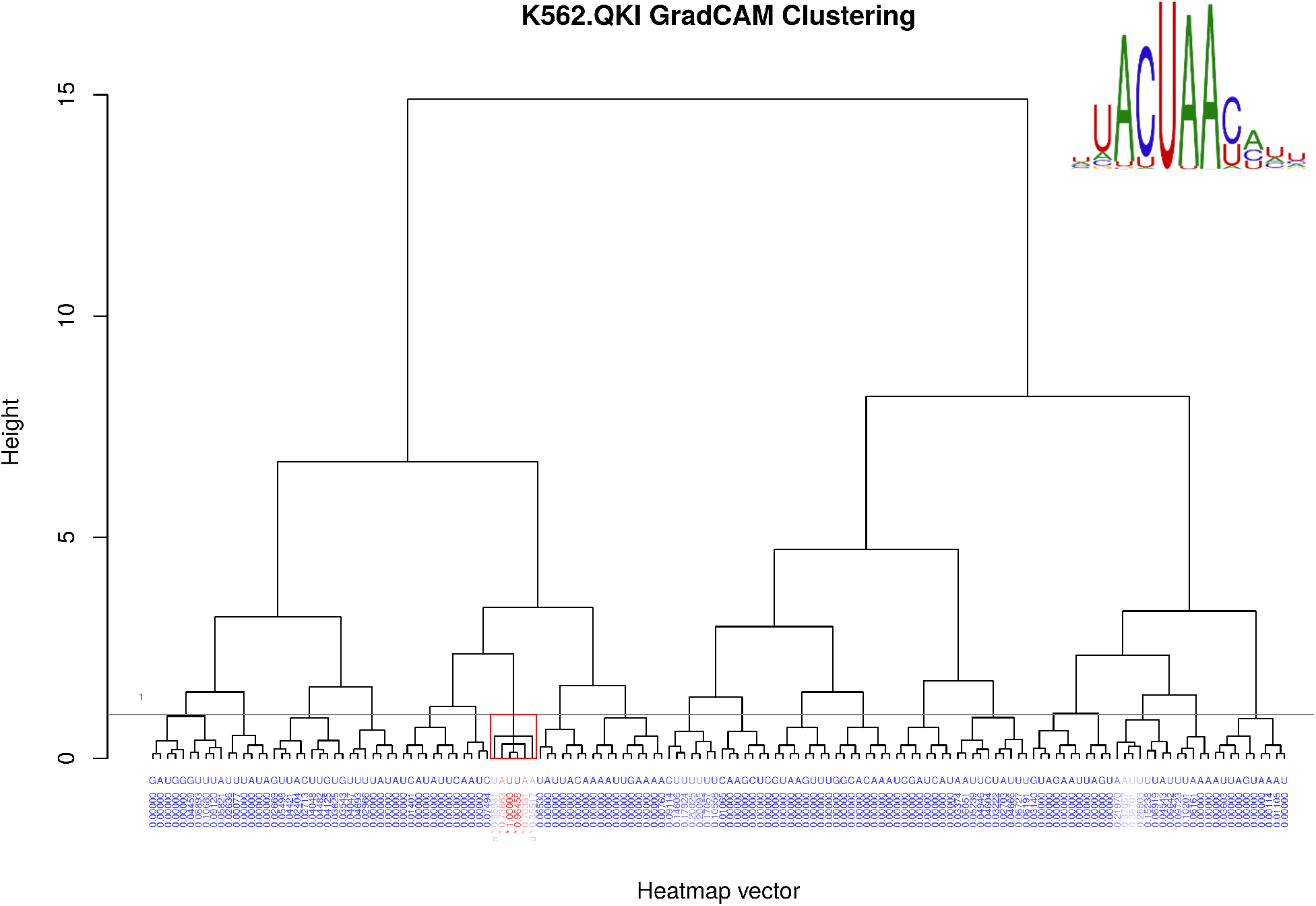
GradCAM heatmap clustering example for QKI. Hierarchical clustering breakdown of a QKI GradCAM heatmap (POLARIS prediction score 0.629), for a region containing chr10:112708640 C→A mutation. This variant was highlighted in our earlier CHD Whole Genome Sequencing study described in Richter, et al. 2020 [27], as its genomic location is near the known (and recurrently hit) CHD gene *SHOC2*. The automatically extracted QKI binding motif UA*UAA (before a less confident U/CAUU) is a good match to the canonical sequence logo pictured in the top right (taken from mCrossBase, as in Figure 7). Clustering parameters used were score (vs. sequence proximity) weight α = 0.9, hierarchical tree cut height h = 1, and motif designation threshold m = 0.5 parameters.

Our weighted hierarchical clustering algorithm can break down GradCAM heatmaps into binding sites and valleys, with flexible motif length and number of motifs per window that better mirror the variability inherent in the diverse binding patterns of different RBPs. This clustering method has difficulty in cases of lower POLARIS binding score, which could perhaps be addressed with hyperparameter tweaks and additional sophistication on distance matrix D to rapidly decay pairwise similarity in beyond certain reasonable motif lengths. Hierarchical tree cut height h = 1 was observed to work well in practice, but could be further optimized based on concrete experiments on RBP motif extraction.

## References

1. Homsy, J., et al., De novo mutations in congenital heart disease with neurodevelopmental and other congenital anomalies. Science, 2015. 350(6265): p. 1262–6.

2. De Rubeis, S., et al., Synaptic, transcriptional and chromatin genes disrupted in autism. Nature, 2014. 515(7526): p. 209–15.

3. Short, P.J., et al., De novo mutations in regulatory elements in neurodevelopmental disorders. Nature, 2018. 555(7698): p. 611–616.

4. Van Nostrand, E.L., et al., Author Correction: A large-scale binding and functional map of human RNA-binding proteins. Nature, 2021. 589(7842): p. E5.

5. Consortium, E.P., An integrated encyclopedia of DNA elements in the human genome. Nature, 2012. 489(7414): p. 57–74.

6. Van Nostrand, E.L., et al., Robust transcriptome-wide discovery of RNA-binding protein binding sites with enhanced CLIP (eCLIP). Nat Methods, 2016. 13(6): p. 508–14.

7. Yang, E.W., et al., Allele-specific binding of RNA-binding proteins reveals functional genetic variants in the RNA. Nat Commun, 2019. 10(1): p. 1338.

8. Zhou, J. and O.G. Troyanskaya, Predicting effects of noncoding variants with deep learning-based sequence model. Nat Methods, 2015. 12(10): p. 931–4.

9. Zhou, J., et al., Whole-genome deep-learning analysis identifies contribution of noncoding mutations to autism risk. Nat Genet, 2019. 51(6): p. 973–980.

10. Quang, D. and X. Xie, DanQ: a hybrid convolutional and recurrent deep neural network for quantifying the function of DNA sequences. Nucleic Acids Res, 2016. 44(11): p. e107.

11. Lee, D., LS-GKM: a new gkm-SVM for large-scale datasets. Bioinformatics, 2016. 32(14): p. 2196–8.

12. Ghandi, M., et al., gkmSVM: an R package for gapped-kmer SVM. Bioinformatics, 2016. 32(14): p. 2205–7.

13. Feng, H., et al., Modeling RNA-Binding Protein Specificity In Vivo by Precisely Registering Protein-RNA Crosslink Sites. Mol Cell, 2019. 74(6): p. 1189–1204 e6.

14. Fukushima, K., Neocognitron: a self organizing neural network model for a mechanism of pattern recognition unaffected by shift in position. Biol Cybern, 1980. 36(4): p. 193–202.

15. LeCun, Y., et al., Backpropagation Applied to Handwritten Zip Code Recognition. Neural Computation, 1989. 1.

16. LeCun Y., H.P., Bottou L., and Bengio Y., Object Recognition with Gradient-Based Learning, in Shape, Contour and Grouping in Computer Vision. J.L.M. David A. Forsyth, Vito di Gesu, Roberto Cipolla, Editor. 1999, Springer. p. 319–345.

17. Yamaguchi, K., Sakamoto, K., Akabane, T., and Fujimoto, Y., A Neural Network for Speaker-Independent Isolated Word Recognition, in First International Conference on Spoken Language Processing (ICSLP 90). 1990: Kobe, Japan.

18. Rumelhart, D.E., G.E. Hinton, and R.J. Williams, Learning representations by back-propagating errors. Nature, 1986. 323(6088): p. 533–536.

19. Pesole, G., et al., Structural and compositional features of untranslated regions of eukaryotic mRNAs. Gene, 1997. 205(1–2): p. 95–102.

20. Shah, A., et al., CLIP Tool Kit (CTK): a flexible and robust pipeline to analyze CLIP sequencing data. Bioinformatics, 2017. 33(4): p. 566–567.

21. Sanchez de Groot, N., et al., RNA structure drives interaction with proteins. Nat Commun, 2019. 10(1): p. 3246.

22. Koo, P.K., et al., Inferring Sequence-Structure Preferences of RNA-Binding Proteins with Convolutional Residual Networks. bioRxiv, 2018: p. 418459.

23. Lorenz, R., et al., ViennaRNA Package 2.0. Algorithms Mol Biol, 2011. 6: p. 26.

24. Kazan, H., et al., RNAcontext: a new method for learning the sequence and structure binding preferences of RNA-binding proteins. PLoS Comput Biol, 2010. 6: p. e1000832.

25. Orenstein, Y., Y. Wang, and B. Berger, RCK: accurate and efficient inference of sequence- and structure-based protein-RNA binding models from RNAcompete data. Bioinformatics, 2016. 32(12): p. i351–i359.

26. Selvaraju, R.R., et al., Grad-CAM: Visual Explanations from Deep Networks via Gradient-Based Localization. 2017 IEEE International Conference on Computer Vision (ICCV), 2016: p. 618–626.

27. Richter, F., et al., Genomic analyses implicate noncoding de novo variants in congenital heart disease. Nat Genet, 2020. 52(8): p. 769–777.

